# P-PPI: accurate prediction of peroxisomal protein-protein interactions (P-PPI) using deep learning-based protein sequence embeddings

**DOI:** 10.1101/2023.06.30.547177

**Authors:** Marco Anteghini, Vitor AP Martins dos Santos, Edoardo Saccenti

## Abstract

Protein-protein interactions (PPIs) are crucial for various biological processes, and their prediction is typically accomplished through experimental methods, which can be time-consuming and costly. Computational methods provide a faster and more cost-effective approach, leveraging protein sequences and other data sources to infer PPIs. Deep learning (DL) approaches have shown promising results in various protein-related tasks, including PPI prediction. However, DL-based embeddings are often not thoroughly compared or evaluated against state-of-the-art tools. Additionally, existing PPI predictors incorporate different types of information beyond protein sequence representation, making it important to assess the effectiveness of DL-based embeddings solely relying on protein sequences. In this work, we benchmark and compare commonly used DL-based embeddings for PPI prediction based solely on protein sequence information. We utilize high-quality training data, including experimentally validated negative interactions from the Negatome database. The best model, obtained through double cross-validation and hyperparameter optimization, is selected and evaluated to predict peroxisomal PPIs. The resulting tool, P-PPI, is further enhanced by combining AlphaFold2-Multimer predictions with the P-PPI model, leveraging DL-based embeddings and protein structure predictions for a comprehensive analysis of peroxisomal PPIs. This integrated approach holds significant potential to advance our understanding of complex protein networks and their functions.

## 1 Introduction

Protein-protein interactions (PPIs) are specific physical contacts that occur between two or more proteins in a cell or living organism. They involve biochemical events where interactions such as electrostatic forces, hydrogen bonding, and the hydrophobic effect take place [1]. PPIs play crucial roles in various biological processes, including signal transduction (a chemical or physical signal is transmitted through a cell as a series of molecular events), enzymatic reactions [2], and gene regulation [3].

Protein-protein interaction can be determined by experimental methods that include techniques such as:

### Yeast two-hybrid (Y2H) assay [4]

This technique involves the use of a yeast strain engineered to express two hybrid proteins - a DNA-binding domain (DBD) and an activation domain (AD) - fused to the proteins of interest. If the proteins interact, the DBD and AD come in close proximity, leading to the activation of a reporter gene;

### Co-immunoprecipitation (Co-IP) [5]

It relies on the immunoprecipitation of a target protein using an antibody specific to that protein. If interacting proteins are present in the sample, they can be co-immunoprecipitated and detected using techniques such as Western blotting or mass spectrometry;

### Affinity purification coupled with mass spectromety (AP-MS) [6]

It is a method that analyzes tagged proteins expressed in cells. It entails purifying the tagged proteins and their associated binding partners through affinity chromatography. Following purification, the protein complexes are eluted and separated using gel electrophoresis or liquid chromatography. Mass spectrometry is then employed to identify the proteins;

### Fluorescence Resonance Energy Transfer (FRET) [7]

Proteins of interest are labeled with fluorophores, and when they interact, energy transfer occurs, leading to a measurable change in fluorescence.

While these experimental methods provide valuable information, they are often time-consuming, expensive, and limited to specific experimental conditions.

Computational methods offer a faster and more cost-effective approach to predict PPIs, in particular if benchmarked against experimental data. These methods leverage various data sources, including protein sequences, structures, evolutionary information, and functional annotations, to infer protein interactions.

The application of deep learning (DL) approaches to encode protein sequences has shown promising results in several tasks such as subcellular [8] and sub-organelle [9–11] classification, protein structure [12, 13] and function prediction [14–17], and protein-protein interactions (PPI) prediction [18, 19].

However, these embeddings are often not compared and evaluated among themselves for highly specific tasks and are not compared against the state-of-the-art tools available. In particular, PPI prediction tools are well-established and already provide highly accurate predictions, but they often utilize different types of information (e.g., structural, evolutionary) or complex algorithms beyond protein sequence representation [20–25, 18]. DL-based embedding predictors offer several advantages. Firstly, they do not require lengthy pre-processing steps, as only the FASTA sequence is necessary for input [9, 10, 17]. This simplifies the prediction process and saves time. Additionally, DL-based embeddings are pre-computed, making the overall procedure computationally efficient [9, 10, 17]. Lastly, these predictors consistently yield highly accurate results [9, 10, 17]. Among the established PPI predictors that incorporate various types of information beyond just the sequence, notable examples include:

### PIPE4 [21]

It is a network-based algorithm that predicts PPIs based on network topology and sequence similarity. It uses a graph-based approach to represent proteins as nodes and their interactions as edges in a network. The algorithm then applies machine learning techniques to predict new interactions based on the properties of the network.

### PEPPI [24]

It is built using a combination of structural similarity, sequence similarity, functional association data, and machine learning-based classification through a naive Bayesian classifier model. The algorithm consists of several independent modules, including a neural network trained on conjoint triads (where seven classes of amino acids are clustered according to their dipoles and volumes of the side chains, and any three continuous amino acids are regarded as a unit, from which 343 features can be extracted) [26], a STRING database lookup module (which checks for known interacting proteins), a threading-based module [27] (a protein modeling method for proteins that have the same fold as proteins of known structures but do not have homologous), and a sequence-based module using the Basic Local Alignment Search Tool BLAST [28, 29]. Scores from each module are transformed into a ratio of likelihood based on pre-trained score probability distributions, and the final likelihood ratio is calculated as the product of likelihood ratios from each independent module.

### PPI-Affinity [30]

It is built using a dataset of protein-protein complexes with known binding affinity (BA) data from the PDBbind database [31]. Moreover, molecular descriptors using ProtDCal were used. The model was trained using Support Vector Machines (SVM) with ensemble learning, optimizing hyperparameters using a grid search, and selecting the best ensemble model based on performance measures.

Moreover, the AlphaFold2 algorithm has demonstrated remarkable accuracy in predicting protein structures, including residues involved in protein-protein interactions [12]. Recent studies applying AlphaFold2-Multimer, have highlighted its suitability and limits for predicting interacting complexes [32, 33, 19]. Again, these complex algorithms use various types of information beyond protein sequence representation. That emphasizes the need to thoroughly evaluate the effectiveness of DL-based embeddings in predicting PPIs, thus solely relying on the protein sequences. It is important to demonstrate the extent to which these embeddings can be employed by the scientific community to achieve quick and precise predictions, ultimately reducing the reliance on extensive initial information and pre-processing steps for the predictor.

It is important to note that computational methods often have PPI training sets which include high quality positive interactions but lack of reliable negative interactions. Instead, negative samples are often generated by randomly pairing proteins that are not known to interact. However, these negative samples may not accurately represent the absence of interactions, leading to potential biases in the prediction models. The Negatome database [1] provides a curated collection of negative PPIs, which can be useful for training and evaluating PPI prediction models.

In this work, we aim to benchmark and compare the most commonly used DL-based embeddings [14, 15, 13] for predicting PPIs solely based on protein sequence information, relying on high quality training data (that includes experimentally validated negative protein-protein interactions).

The best model, obtained through double cross-validation (DCV) [34] and hyperparameter optimization, is selected and further evaluated to predict peroxisomal protein-protein interactions. The resulting tool is referred to as Peroxisomal Protein-Protein Interaction (P-PPI). Additionally, this study explores the complementary use of the P-PPI tool and AlphaFold2 [12]. By combining AlphaFold2-Multimer predictions with P-PPI, we aim to enhance the assessment of protein-protein interactions by incorporating structural insights. This integrated approach leverages the strengths of both computational methods, employing DL-based embeddings and precise protein structure predictions to enable a comprehensive and accurate analysis of peroxisomal protein-protein interactions. The integration of these approaches has significant potential to help us advance our understanding of complex protein networks and their functions.

## 2 Methods

### 2.1 Data set

#### 2.1.1 Training and validation set

The PPIs are retrieved from the DIP database at https://dip.doe-mbi.ucla.edu/dip/Main.cgi(human dataset - 2017). The positive dataset contains 5970 encoded PPIs. The non-PPIs are retrieved from NEGATOME 2.0 at http://mips.helmholtz-muenchen.de/proj/ppi/negatome/manual.html. The negative dataset contains 2087 encoded non-PPIs. As a representative and more balanced sample, we retrieved positive and negative training sets of size 1500 and a validation set of size 1000.

#### 2.1.2 Peroxisomal PPI data set

We selected non overlapping protein-protein interactions (from the protein left out from the DIP data set) where at least one of the two proteins was a peroxisomal protein. In total we obtained 77 peroxisomal interactions. 60 where used for fine-tuning the model. 17 where used as test set.

### 2.2 Deep Learning Protein Sequence Embeddings

We considered three recently proposed methods for the embedding of protein sequences based on deep-learning approaches:

#### Unified Representation (UniRep)

UniRep [14] employs a recurrent neural network architecture with 1900 hidden units to capture chemical, biological, and evolutionary information from protein sequences. It utilizes a hidden state vector that is iteratively updated based on the previous state. This allows the model to learn by predicting the next amino acid in the sequence based on the preceding context. UniRep generates protein sequence embeddings of length 64, 256, or 1900 units, depending on the chosen neural network architecture. In this study, we utilized the 1900-unit embedding (average final hidden array). For a detailed explanation on how to obtain the UniRep embedding, please refer to the specific GitHub repository: https://github.com/churchlab/UniRep.

#### Sequence-to-Vector (SeqVec)

The Sequence-to-Vector embedding (SeqVec) [15] adopts a natural language processing approach, treating amino acids as words and proteins as sentences, to capture biophysical information from protein sequences. SeqVec is obtained by training ELMo [35], a deep contextualized word representation that models complex characteristics of word use and their variations across contexts, using a 2-layer bidirectional LSTM [36] backbone. In this case, ELMo is pretrained on a large text corpus, specifically UniRef50 [37]. SeqVec can generate embeddings at both the per-residue (word-level) and per-protein (sentence-level) levels. The per-residue level embedding allows prediction of secondary structure or intrinsically disordered regions, while the per-protein level embedding enables prediction of subcellular localization and differentiation between membrane-bound and water-soluble proteins [15]. In this study, we utilized the per-protein level representation, where the protein sequence is represented by a 1024-unit embedding. For detailed instructions on retrieving the SeqVec embedding, please refer to the specific GitHub repository: https://github.com/mheinzinger/SeqVec.

#### Evolutionary Scale Modelling - 1b (ESM-1b)

ESM-1b is trained on 250 million sequences from the UniParc database [38] and utilizes a deep transformer architecture [39, 40], a powerful model for representation learning and generative modeling in natural language processing (NLP). The transformer architecture in ESM-1b provides contextual information for each amino acid (word) in the sequence (sentence) by comparing it to every other amino acid (word) in the sequence, including itself, using self-attention blocks. These blocks consist of three steps: 1) Computing dot product similarity and alignment scores; 2) Normalizing the scores and weighting the embeddings; 3) Re-weighing the original embeddings based on the scores. In ESM-1b, the transformer processes inputs through blocks that alternate self-attention with feed-forward connections. Since it is trained on proteins, the self-attention blocks capture pairwise interactions between all positions in the sequence, representing residue-residue interactions. ESM-1b is trained using masked language modeling objective, which requires the model to predict masked parts of the sequence by understanding dependencies between the masked site and the unmasked parts. The model is optimized with hyperparameters to train a 650 million-parameter (33 layers) model on the UR50/S dataset, resulting in the ESM-1b Transformer [13]. The resulting ESM-1b vector has a length of 1280 units.

### 2.3 Step Forward Feature Selection

As reported in our previous study where we analysed DL-based sequence embedings, also here we adopted a Step Forward Feature Selection (SFFS) approach [9]. SFFS was used to select the best combination of features (predictors) that is, protein encodings or embeddings to be used as input for classification algorithms [41]. It is a wrapper method that evaluates subsets of variables, in our case, combinations of protein embeddings. It starts with the evaluation of each individual encoding, and selects that which results in the best performing selected algorithm model. Next, it proceeds by iteratively adding one embedding to the current best performing features and evaluating the performance of the classification. The procedure is halted when performance worsens and the best combination of embeddings is retained.

### 2.4 Logistic Regression (LR)

We used a penalised implementation of multivariable logistic regression [42]. Penalized multivariable logistic regression is a technique used to handle high-dimensional data with multicollinearity and overfitting issues. It incorporates a penalty term to control model complexity and select relevant predictors. One commonly used penalized approach is ridge regression, which adds a squared L2 penalty to the objective function. Another popular penalized method is lasso regression, which adds an L1 penalty to the objective function. Ridge regression shrinks coefficients towards zero, lasso regression performs variable selection, and elastic net combines both methods. Penalized logistic regression improves model generalizability, mitigates overfitting, and enhances interpretability [42, 43].

### 2.5 Model Calibration and Validation

We employed a double cross-validation (DCV) technique [34, 44] for two main purposes: (i) optimizing the hyperparameters of the classification algorithms used, enabling model calibration, and (ii) obtaining unbiased estimates of prediction errors when applying the model to new cases within the dataset population. This approach is particularly suitable for small datasets.

The DCV strategy involves two nested cross-validation loops. In the outer loop, the data is initially divided into *k* folds, with one fold serving as the validation set while the remaining *k* − 1 folds are used for model calibration. The inner loop is applied to the calibration set, further splitting it into training and test sets using a *k*-fold division. In our study, we utilized 5 folds for both the inner and outer loops.

The inner loop is responsible for optimizing the hyperparameters of the different classification algorithms. A (hyper)grid search is performed, evaluating the average classification score across the folds for each set of hyperparameters. The hyperparameters corresponding to the best classification score are then used to fit a classification model, and its quality is assessed on the validation set. This ensures unbiased evaluation of the model since the validation data was not used during the training of the classification model.

Additionally, we incorporated a step-forward feature selection procedure, as described in Section 2.3, within the calibration loop. This means that model calibration also involved selecting the best combination of protein sequence encodings and embeddings, based on their predictive ability.

Given the imbalance between the two classes of proteins, different weights were applied to each class. These class weights were considered metaparameters and were optimized within the inner calibration loop.

### 2.6 AlphaFold2-Multimer

The AlphaFold system [12], described in Jumer *et al*. (2021), is a protein structure prediction method that integrates various sources of information such as the amino acid sequence, multiple sequence alignments (MSA) [45, 46], and homologous structures. It utilizes a neural network called Evoformer, which incorporates a neural representation of the MSA and pairwise relations between amino acids [12].

The pairwise representation provides information about the relative positions of amino acids in the chain and is used to estimate the relative distances between them. The first row of the MSA embedding, together with the pair embedding, is used to generate the final structure prediction. The model is trained end-to-end (all of the parameters are trained jointly), with gradients propagating from the predicted structure through the entire network [12].

AlphaFold2-Multimer is an extension of the original AlphaFold system that specifically addresses the prediction of protein complexes with multiple chains [32]. One of the key challenges in modeling protein complexes is accounting for permutation symmetry [32, 47]. Since a protein sequence can appear multiple times in a complex, the model cannot rely on predicting the chains in a predetermined order [47]. AlphaFold2-Multimer incorporates an optimization process to identify the optimal permutation of predicted homomer chains that aligns with the ground truth. This process utilizes a simple heuristic that greedily seeks a suitable ordering for the chains [33].

The approach for constructing multiple sequence alignments (MSAs) in AlphaFold2-Multimer [32] follows a similar methodology to that employed in AlphaFold [45, 46, 12]. For homomeric complexes, the MSA is replicated X times, where X denotes the number of chain repeats, and aligned in a left-to-right stacking manner. In the case of heteromeric complexes, the sequences between the MSAs of each chain are paired whenever possible, utilizing UniProt species annotation [48, 49]. The ranking of candidate rows for each chain is determined based on their similarity to the respective target sequence, and pairs with the same rank are concatenated. If partial alignments exist, with sequence pairings between certain chains but not others, gaps are introduced between the paired sequences as padding. The remaining unpaired MSA sequences for each chain are stacked in a block-diagonal fashion below the set of paired sequences, with padding applied on the off-diagonal regions. During training, sampling of the MSA cluster centers is biased to ensure an equal probability of sampling each chain’s unpaired MSA, regardless of the number of sequences it contains. The sampling of paired and unpaired sequences is unbiased and proportional to their relative occurrence. During testing, unbiased MSA sampling is performed to enhance performance slightly, as it introduces greater diversity across recycling iterations [32].

AlphaFold2-Multimer system employs a cropping procedure aimed at maximizing chain coverage and crop diversity, while also ensuring a balanced representation of interface and non-interface regions [32]. Due to memory and compute limitations, the number of residues that can be trained on is restricted, and thus the cropping process involves selecting contiguous blocks of residues up to a maximum length of 384 residues. The cropping procedure is specifically designed to involve multiple chains within a complex and give priority to binding interfaces between the chains, as these interfaces play a critical role in accurately modeling protein complexes [32].

The training protocol for AlphaFold2-Multimer closely adheres to that of AlphaFold. The training dataset comprises protein structures sourced from the Protein Data Bank (PDB) up to a maximum release date of 2018-04-30 [50, 32].

#### 2.6.1 AlphaFold metrics

##### The Local Distance Difference Test (lDDT) [51]

It is a superposition-free score that evaluates local distance differences of atoms in a model, including validation of stereochemical plausibility [51]. lDDT reports the domain accuracy without requiring a domain segmentation of chain structures [32]. The distances are either computed between all heavy atoms (lDDT) or only the C*α* atoms to measure the backbone accuracy (lDDT-C*α*).

##### Predicted aligned error (PAE) [32]

The position error of AlphaFold is assessed for each pair of residues, providing an estimation of the distance error between the predicted and true structures when aligned on specific residues. The range of values for the position error is typically from 0 to 35 Angstroms. This information is commonly visualized as a heatmap, where the residue numbers are displayed along the vertical and horizontal axes, and the color at each pixel represents the position error value (PAE) for the corresponding pair of residues. When the relative position of two domains is accurately predicted, the PAE values will be low (typically less than 5 Angstroms) for residue pairs with one residue in each domain.

### 2.7 Metrics

The evaluation metrics used for assessing the performance of the models in binary classification (true vs. false interaction) were accuracy (ACC), F1 score [52], Matthews correlation coefficient (MCC) [53], and the area under the curve (AUC) of the receiver operating characteristic (ROC). These metrics are calculated based on the number of true positives (*TP*), false positives (*FP*), true negatives (*TN*), and false negatives (*FN*) using the following formulas:

True positive rate (TPR) is defined as

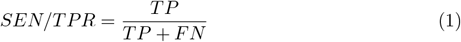

False positive rate (FPR) is defined as

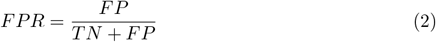

Accuracy (*ACC*) is defined as

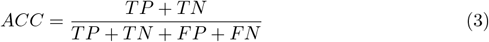

*F*_1_ score [52] is defined as

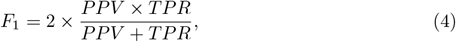

where *PPV* is the positive predicted value (or precision)

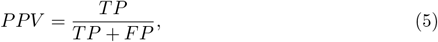

The *F*_1_ score is the harmonic mean of recall and precision and varies between 0, if the precision or the recall is 0, and 1 indicating perfect precision and recall.

Matthews correlation coefficient (MCC) [53] is defined as

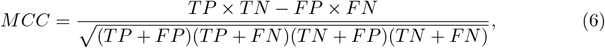

*MCC* is the correlation coefficient between the true ad predicted class: it is bound between −1 (total disagreement between prediction and observation) and +1 (perfect prediction); 0 indicates no better than random prediction. The *MCC* is appropriate also in presence of class unbalance [54].

The area under the curve (AUC) of the receiver operating characteristic (ROC) curve which plots the true positive proportion or the Sensitivity against the Specificity, is defined as

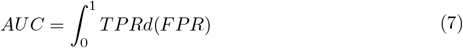

The AUC analysis enables the evaluation of the performance of a binary classifier system according to the variation of the discrimination threshold. A perfect prediction has an AUC score of 1.0 while an AUC of 0.5 indicates randomness [55].

### 2.8 Approach

We evaluated three different DL-based embeddings, SeqVec, UniRep, ESM-b1 for correctly predicting PPIs. To do so we performed 5-fold Double Cross Validation (DCV) with a Logistic Regression (LR) classifier. The best model was then tested against the validation set.

## 3 Results

### 3.1 P-PPI pipeline overview

The Figure 1 provides an overview of the pipeline developed in this study, showcasing the integration of the P-PPI model and an optional analysis using AlphaFold2-Multimer.

**Figure 1.**
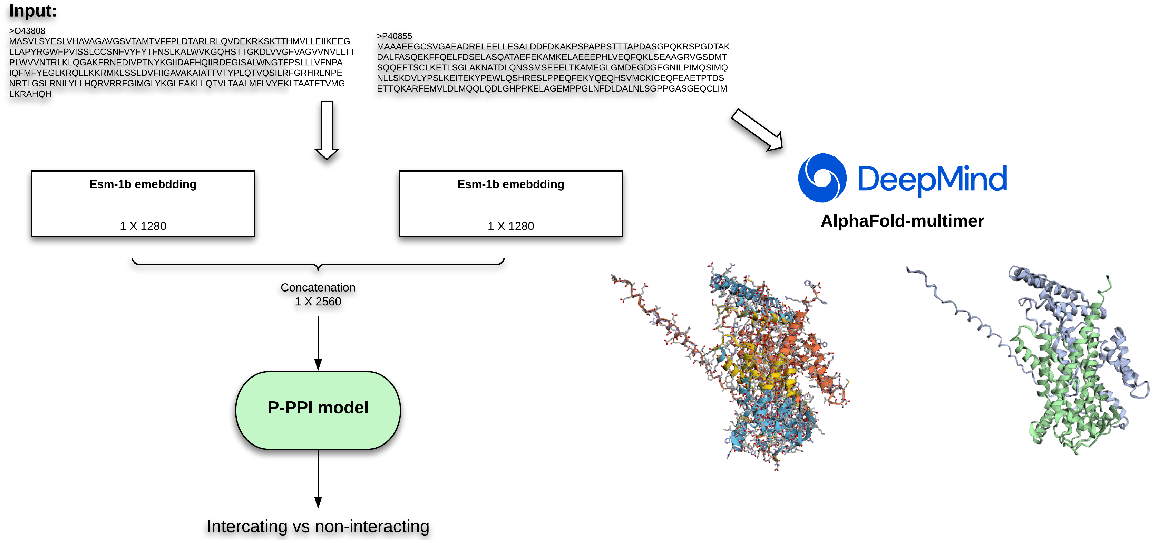
Overview of the P-PPI pipeline workflow: The protein sequences of the two putative interacting proteins (in FASTA format) enter the pipeline and are initially encoded using the ESM-1b embedding. The resulting embeddings are then concatenated to represent the protein interaction. The P-PPI model, consisting of the ESM-1b embedding and Logistic Regression, predicts whether the interaction is true or false. Additionally, as an optional step, the model can be validated against AlphaFold2-Multimer to assess the quality of complexes that involve the two interacting proteins. A robust and precise model indicates the reliability of the predicted interaction.

### 3.2 P-PPI model performances with different protein embed-dings

The classification performance of various embeddings, including their concatenation, in distinguishing true and false interactions was evaluated using a step-forward feature selection approach. The results are presented in Table 1. Esm-1b demonstrated superior performance compared to SeqVec and UniRep, achieving accuracy scores of 0.8703, 0.8656, and 0.8654, respectively. The performance of the concatenated Esm-1b and SeqVec embeddings was found to be very similar to the performance of the Esm-1b single embedding (0.8703 vs 0.8716). Consequently, we utilized the single embedding representation (Esm-1b) for the final P-PPI model. Overall, the performances of all embeddings are very close.

**Table 1.** Results obtained using SeqVec, UniRep, and ESM-b1 embeddings are reported in terms of accuracy (ACC), Matthew’s Correlation Coefficient (MCC), *F*_1_ macro score (*F*_1_), and area under the curve (AUC) for both training and validation sets. The best performing embedding is highlighted in bold. The used classifier is Logistic Regression.

| | ACC | | MCC | | $F_1$ | | AUC | |
| --- | --- | --- | --- | --- | --- | --- | --- | --- |
|  | CV | Val. | CV | Val. | CV | Val. | CV | Val. |
| SeqVec | 0.8656 ± 0.0024 | 0.6760 | 0.6114 ± 0.0104 | 0.3573 | 0.7834 ± 0.0058 | 0.6731 | 0.7453 ± 0.0049 | 0.6760 |
| UniRep | 0.8654 ± 0.0021 | 0.6560 | 0.6100 ± 0.0120 | 0.3144 | 0.7840 ± 0.0043 | 0.6548 | 0.7467 ± 0.0017 | 0.6560 |
| **ESM-1b** | 0.8703 ± 0.0144 | 0.6840 | 0.6277 ± 0.0096 | 0.3713 | 0.7883 ± 0.0033 | 0.6827 | 0.7475 ± 0.0102 | 0.6840 |
| UniRep-SeqVec | 0.8579 ± 0.0022 | 0.6630 | 0.5906 ± 0.0133 | 0.3400 | 0.7591 ± 0.0011 | 0.6631 | 0.7179 ± 0.0032 | 0.6630 |
| **ESM-1b-SeqVec** | 0.8716 ± 0.0073 | 0.6860 | 0.6319 ± 0.0059 | 0.3773 | 0.7910 ± 0.0061 | 0.6913 | 0.7502 ± 0.0019 | 0.6860 |
| ESM-1b-UniRep | 0.8556 ± 0.0080 | 0.6560 | 0.5819 ± 0.0049 | 0.3115 | 0.7543 ± 0.0097 | 0.6522 | 0.7143 ± 0.0029 | 0.6548 |

### 3.3 Performances of the peroxisomal proteins fine-tuned model

We fine-tuned the P-PPI model by retraining it on a data set consisting of 60 peroxisomal protein-protein interactions, which was non-overlapping with the original training set. Subsequently, we evaluated the model’s performance on a separate test set comprising 17 peroxisomal interactions. The fine-tuning process resulted in significant improvements, with an accuracy of 0.9118. Detailed results can be found in Table 2.

**Table 2.** Performances of the Logistic Regression model with the ESM-1b embedding against the peroxisomal test set (17 interactions) after fine-tuning. Results are reported in terms of accuracy (ACC), Matthew’s Correlation Coefficient (MCC) F1 macro score (F1) and area under the curve (AUC)

| Metric | Value |
| --- | --- |
| ACC | 0.9118 |
| $F_1$ | 0.9189 |
| MCC | 0.8367 |
| AUC | 0.9250 |

### 3.4 AlphaFold2-Multimer model use case

As a proof of concept, we selected a positively predicted interaction from the P-PPI model and tested it against the AlphaFold2-Multimer algorithm to generate the structure of the interacting complex. Among the 17 PPIs from the peroxisomal test set, we chose a specific interaction (UniProt IDs: P40855-O43808). This interaction comes with 4 experimental validations (e.g. Co-IP); for details we invite to check the Uniprot ID O43808 at https://www.uniprot.org/.

To generate the structure of the interacting complex, we inputted the FASTA sequences of the interacting proteins into the AlphaFold Google Colab script, available at https://colab.research.google.com/github/deepmind/alphafold/blob/main/notebooks/AlphaFold.ipynb. The results of the structure prediction are presented in Figure 2 and the corresponding evaluation metrics are shown in Figure 3.

**Figure 2.**
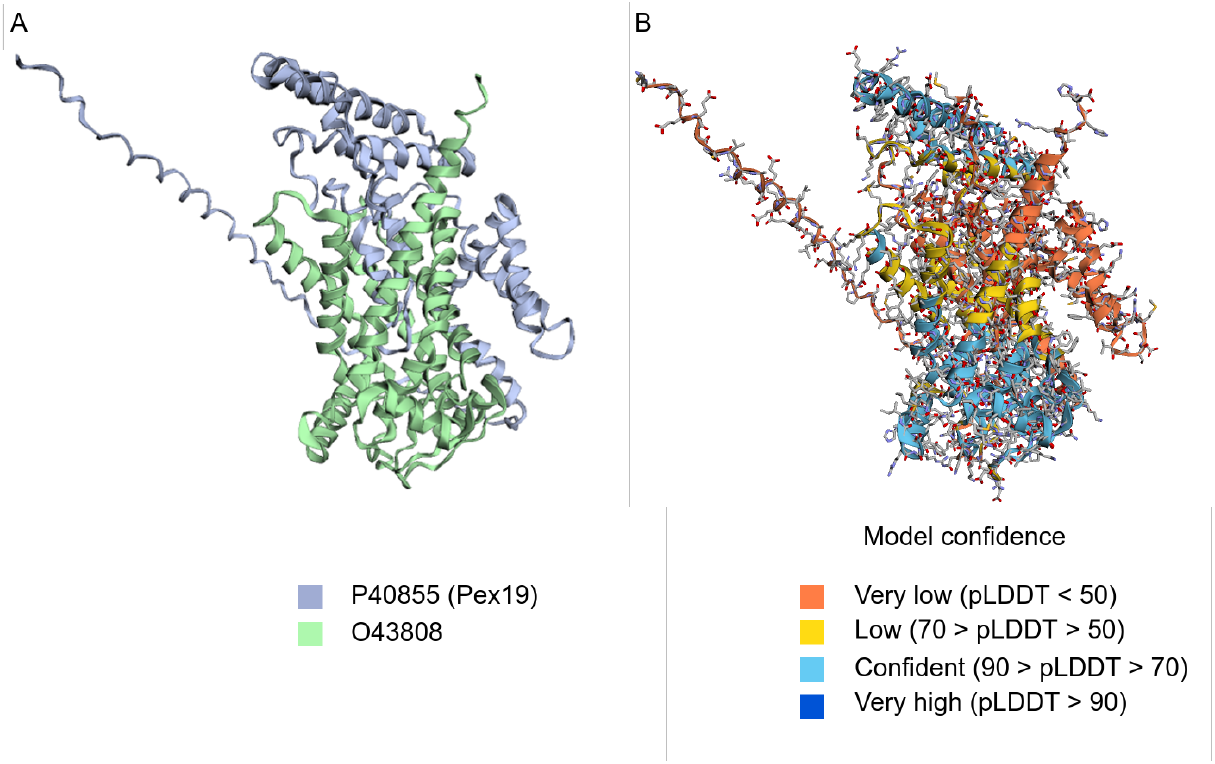
Predicted interacting complex (UniProt IDs: P40855-O43808). Subfigure A, highlights the two proteins in grey (P40855) and light green (O43808). Subfigure B shows the different protions of the complex colered by model confidence levelm expressed with predicted Local Distance Difference Test (pLDDT)

**Figure 3.**
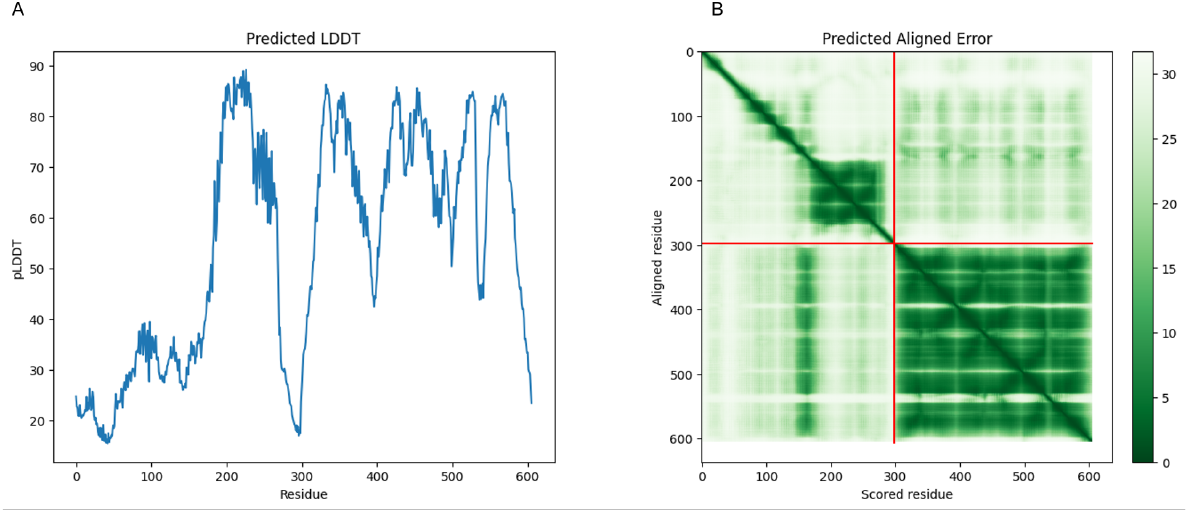
Visualization of the AlphaFold metrics for model evaluation. Subfigure A shows the performances in terms of predicted Local Distance Difference Test (pLDDT) from residue 0 to residue 606 of the modelled complex. Subfigure B shows the performances in terms of Predicted Aligned Error (PAE) as an heatmap from residue 0 to residue 606 of the complex.

## 4 Discussion and Conclusion

The results obtained from our PPI prediction method, as demonstrated in Tables 1 and 2, show promising outcomes and highlight the potential of employing pre-trained deep learning models for protein sequence embedding in the prediction of protein-protein interactions (PPIs). These findings hold true not only for PPIs in general but also for peroxisomal PPIs specifically, thus we implemented the P-PPI model, which accurately predicts peroxisomal protein-protein interactions with an accuracy of 91%.

This opens up possibilities for further analysis and improvements in the field. Our approach involved using the Esm-1b embedding, which proved to be the most effective. We incorporated Esm-1b in the final model, along with the fine-tuned version, resulting in accurate prediction of peroxisomal PPI.

An additional computational validation was performed using AlphaFold2, which provided additional confidence in our predictions as presented in Figures 2 and 3.

It is important to emphasise that our training data set relies on experimentally validated positive and negative interactions. PPI predictors are often built using negative interactions, where the non-interacting proteins are selected from different subcellular locations. These interactions do not occur in a living cell but can be verified in vitro, thus adding noise to the predictive models. The models can inadvertently learn different subcellular locations instead of true and false interactions. Moreover, due to the nature of peroxisomes and their high interconnection with several organelles, we believe it is more relevant to focus on the interactions rather than the subcellular locations.

Moving forward, there are several avenues for future work. Firstly, we plan to test additional deep learning embeddings, such as the novel ESM2, to explore their performance in PPI prediction. Secondly, we aim to create a negative set of non-PPIs based on proteins from different subcellular locations, allowing for a more comprehensive comparison and providing experimental validation to our theory of preferring experimentally validated negative interactions over non-PPIs based on proteins from different subcellular locations. It is also important to consider comparing the Negatome data set with other negative datasets to understand the limitations of alternative data sources in capturing negative interactions.

Furthermore, we intend to compare our predictor against state-of-the-art methods and train the model on PPIs from other organisms. Lastly, we will suggest candidate PPIs and validate the predictions experimentally.

## Notes

### Competing Interest Statement

The authors have declared no competing interest.

